# Paternal transmission of the APPPS1 transgene is associated with increased amyloid pathology and a small Thy1.2–APP methylation difference

**DOI:** 10.64898/2026.11.22.768387

**Authors:** Mirjam Brackhan, Iván Eiriz, Thomas Brüning, Skarphéðinn Halldórsson, Jacob Gildenblat, Jens Pahnke

**Author notes:** Universidad Alfonso X el Sabio, Facultad de Veterinaria, Avenida de la Universidad 1, 28691 Villanueva de la Cañada, Madrid, Spain (Mirjam Brackhan).

## Abstract

Maternal transmission of the APP transgene lowers amyloid burden in 5xFAD mice. We examined whether APPPS1-21 shows a similar parent-of-origin association and whether Thy1.2 methylation contributes. F1 offspring inheriting the transgene from the mother had less A*β*42, fewer plaques, less gliosis, and less human APP at intermediate ages. The A*β*42 difference was not detected in the F2 cohort at 100 days. Non-transgenic off-spring of APPPS1 mothers did not differ in glial counts, and serum anti-A*β* antibody levels were not higher at 40 days. In an exploratory methylation analysis of 32 brains, parent-of-origin-associated differences were sex-specific and enriched on chromosome X, not chromosome 2. The Thy1.2–APP cassette showed an approximately three-percentage-point paternal increase; the PSEN1 cassette did not. These findings establish an association with parental transmission but do not show that methylation causes the pathology differences.

**HIGHLIGHTS:**

- Maternal transmission is associated with lower amyloid pathology
- The amyloid-beta 42 difference was not detected in the second generation
- Amyloid precursor protein cassette methylation is higher with paternal transmission
- The methylation difference does not establish a causal mechanism

**IN BRIEF:** We report lower amyloid pathology after F0 maternal APPPS1 transmission to F1. The A*β*42 difference was not detected in the F2 cohort. The small Thy1.2–APP methylation difference does not establish a causal mechanism.

## INTRODUCTION

Amyloid-*β* (A*β*), a cleavage product of the amyloid precursor protein (APP), accumulates in the brain in Alzheimer’s disease, and APP-transgenic mice remain the main experimental system for studying that accumulation.^1–3^ APPPS1-21 mice co-express human APP with the Swedish mutation and presenilin-1 with the L166P mutation, each under a mouse Thy1.2 promoter. The two constructs were co-injected, the line is maintained hemizygous on C57BL/6J, and the insertion was mapped genetically to the distal arm of chromosome 2.^4^ Standard breeding transmits the transgene from hemizygous males to non-transgenic females.

Two recent papers in *Neuron* ^5,6^ showed that this breeding direction is not neutral in 5xFAD mice. Paternal transmission of the 5xFAD transgene roughly doubled plaque load relative to maternal transmission across colonies; the difference was independent of sex, maternal care, and gestational environment.^6^ An independent 5xFAD founder line crossed with APOE4 showed even stronger suppression of parenchymal plaques and cerebral amyloid angiopathy after maternal inheritance.^5^ Sasmita et al. proposed genomic imprinting of the Thy1.2-driven transgene as a possible explanation and identified the shared Thy1.2 promoter as a candidate; the Saadi et al. report corroborated the inheritance-associated pathology in a different founder line but did not test this methylation mechanism.

In our APPPS1-21 colony, offspring of APPPS1 mothers had less A*β* than offspring of APPPS1 fathers. We quantify this association across plaque stages, assess whether the A*β*42 difference is detected in F2, and examine alternative explanations. We also analyze nanopore methylation in F0 and F1 brains to test the proposed Thy1.2-cassette mechanism, distinguishing it from methylation of endogenous Thy1.

## RESULTS

### Maternal transmission lowers A*β*, plaques, and gliosis in F1 mice

Hemizygous APPPS1 males or females were crossed to non-transgenic C57BL/6J partners to produce F1 offspring that inherited the transgene from the father or the mother (Figure 1). All F0 breeders themselves had a paternally inherited transgene. Soluble and insoluble A*β*42 were measured at 50, 100, 150, and 200 days.

**Figure 1:**
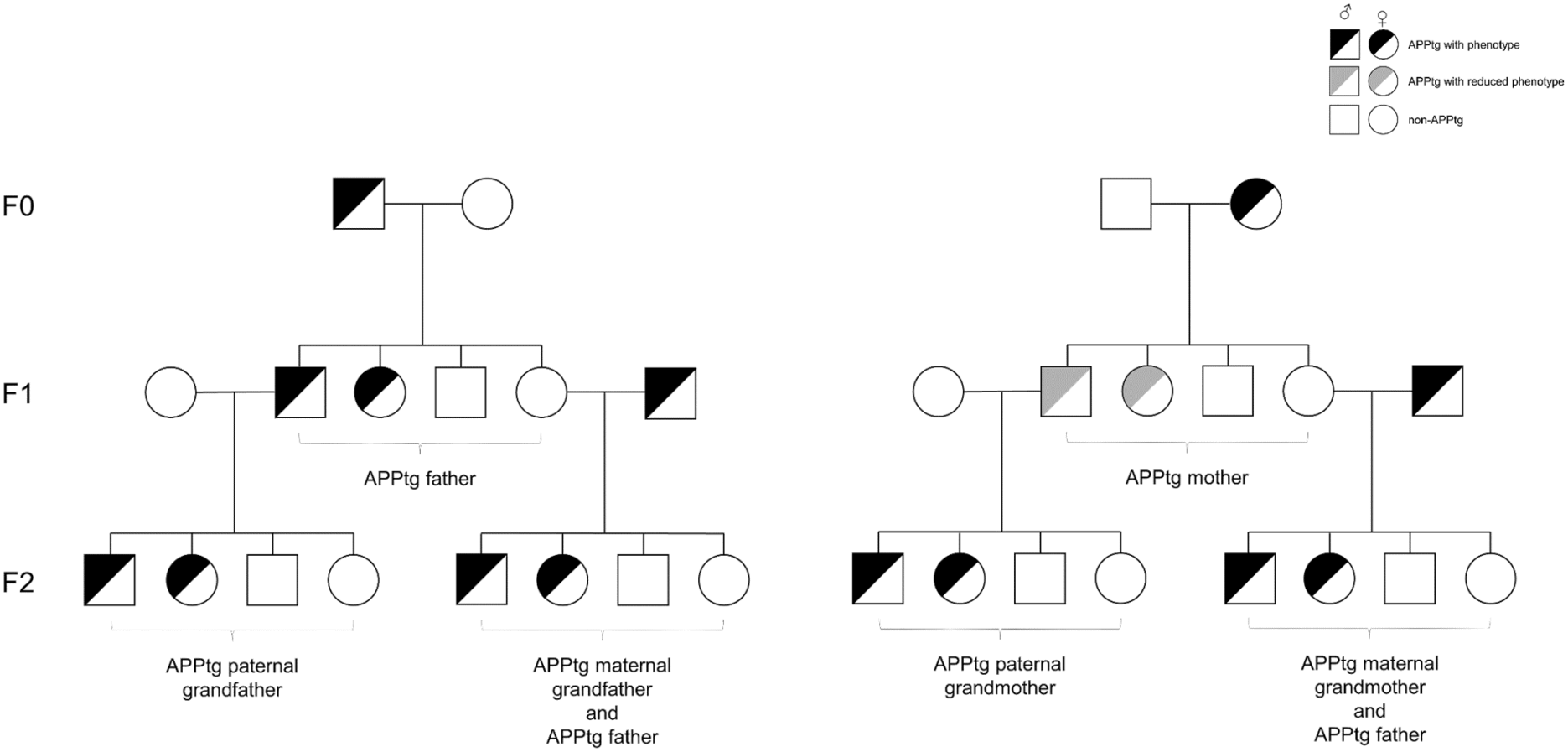
Breeding design. Hemizygous APPPS1 mice (black) that inherited the trans-gene from the father were compared with hemizygous offspring (gray) that inherited it from the mother. F2 crosses test whether the F1 A*β*42 difference is detected after the trans-gene passes through an F1 father and whether non-transgenic daughters of APPPS1 mothers affect A*β*42 in offspring that receive the transgene from the father. Squares, males; circles, females; open symbols, non-transgenic.

Maternal transmission was associated with lower soluble A*β*42 at every age, with reductions of 30.8% at 50 days (*p <* 0.0001), 48.3% at 100 days (*p* = 0.0006), 22.8% at 150 days (*p* = 0.0147), and 34.6% at 200 days (*p* = 0.0005). The insoluble fraction was reduced further: 76.3%, 66.2%, 34.7%, and 46.0% at the same ages (*p <* 0.0001, *p <* 0.0001, *p* = 0.0011, and *p <* 0.0001; Figure 2). Cortical A*β* plaque coverage fell by 65.6%, 36.9%, and 42.3% at 100, 150, and 200 days (each *p <* 0.0001). Total plaque number fell by 60.0%, 32.9%, and 42.3% (*p <* 0.0001, *p <* 0.0001, and *p* = 0.0004); small-plaque counts fell by 57.1%, 30.0%, and 30.2% (*p <* 0.0001, *p <* 0.0001, and *p* = 0.0135). Medium- and large-plaque counts were also lower at each age (all *p <* 0.0001; Figure 3). Activated microglia were fewer by 33.4%, 24.4%, and 29.2% (all *p <* 0.0001), and astrocytes by 50.9%, 25.1%, and 19.8% (*p <* 0.0001, *p <* 0.0001, and *p* = 0.0002; Figure 4).

**Figure 2:**
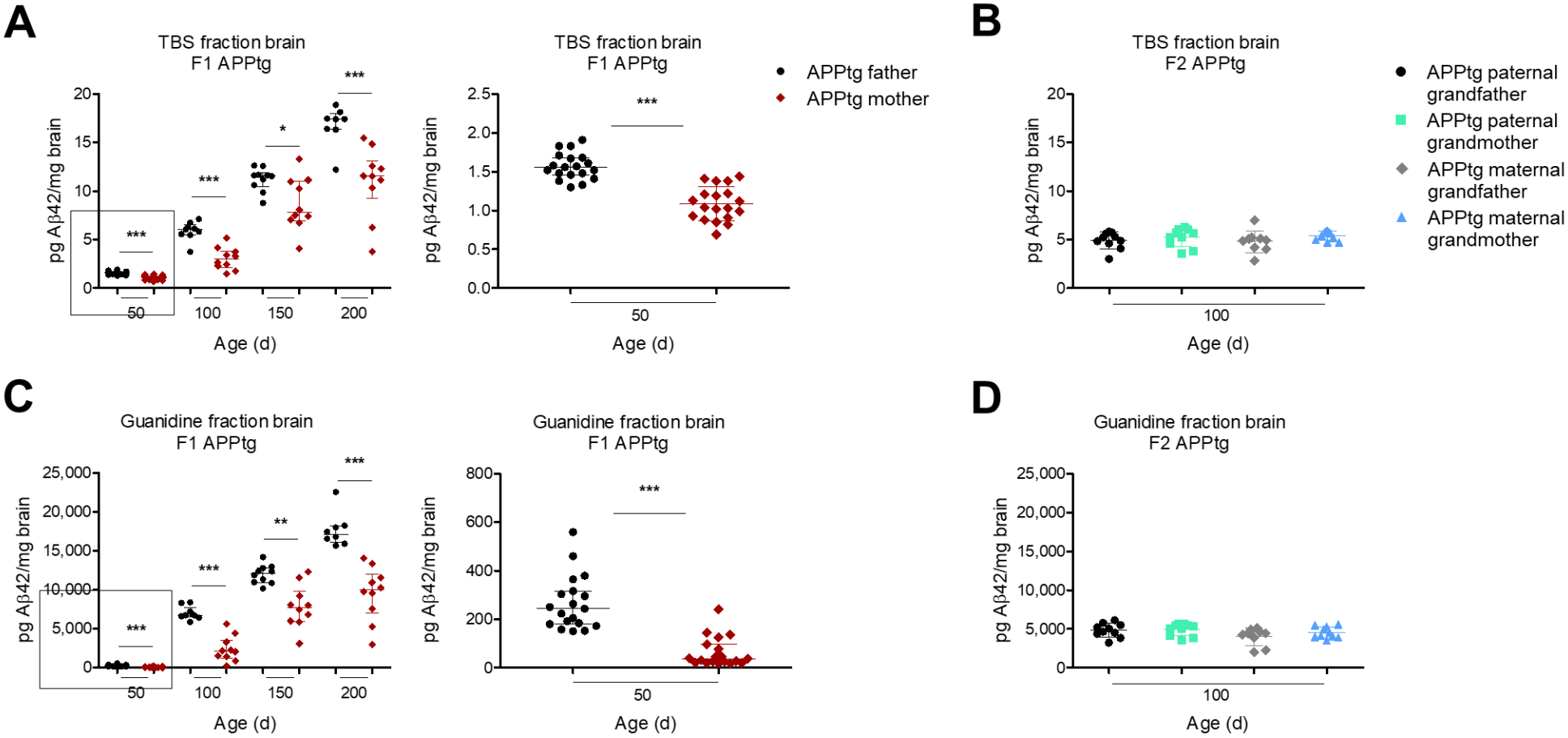
Cerebral A*β*42 is lower after maternal transmission in F1; a difference was not detected in F2. (A, C) Soluble and insoluble A*β*42 in F1 APPPS1 mice from APPPS1 fathers or mothers at 50, 100, 150, and 200 days. Sample sizes: *n* = 19 per group at 50 days; *n* = 9 (father) and 10 (mother) at 100 days; *n* = 10 per group at 150 days; *n* = 8 (father) and 10 (mother) at 200 days. Two-tailed Mann–Whitney tests. Medians and interquartile ranges. The middle plots show expanded views of the 50-day F1 comparisons included in (A) and (C). (B, D) The same fractions in 100-day-old F2 mice (*n* = 10 per group, except *n* = 9 for the maternal-grandfather class). Planned within-branch paternal-grandfather versus paternal-grandmother and maternal-grandfather versus maternal-grandmother comparisons used unpaired two-tailed *t* tests; neither comparison was significant. Means and standard deviations. Values are normalized to brain weight.

**Figure 3:**
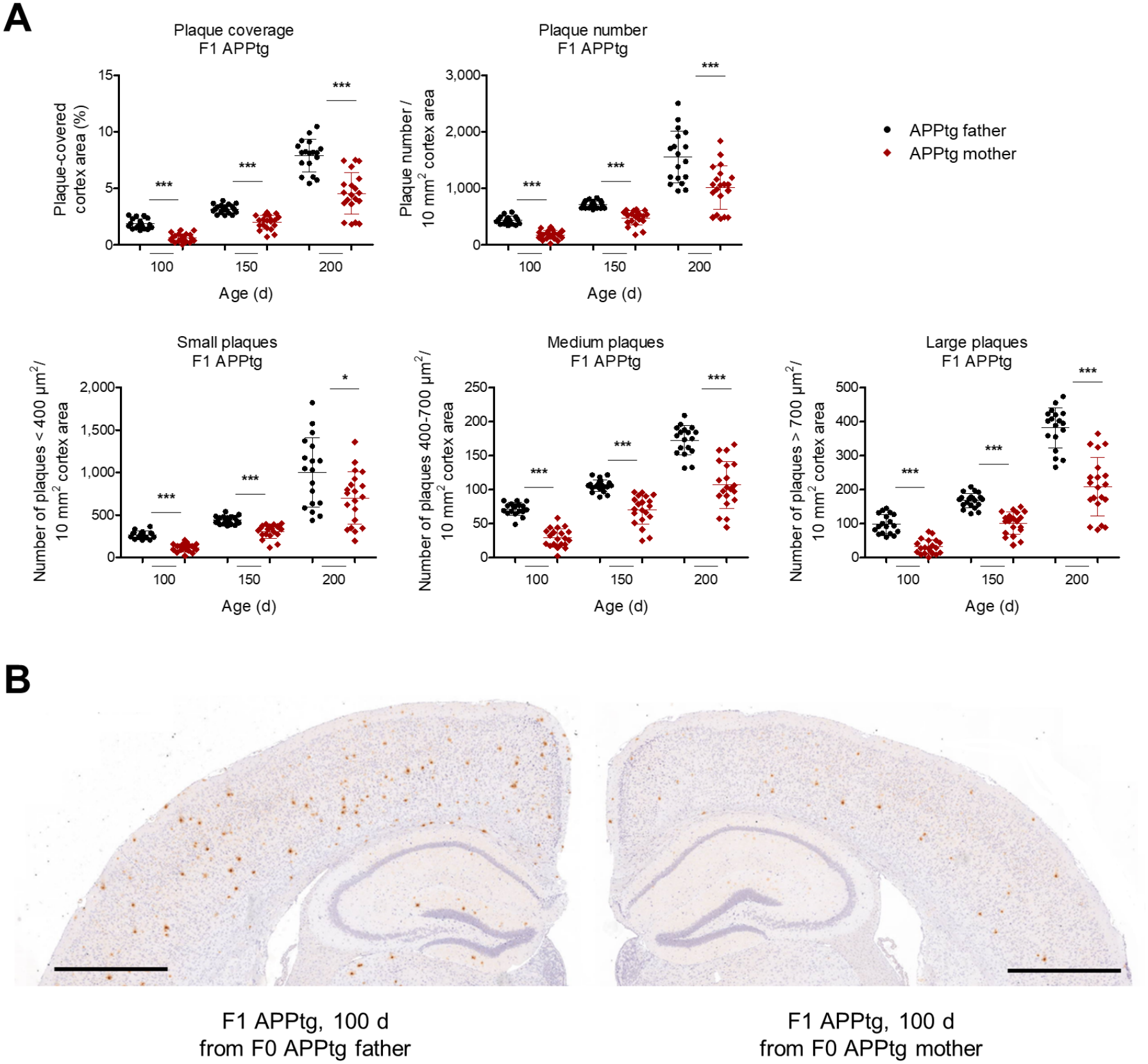
Plaque load is lower after maternal transmission. (A) Cortical A*β* plaque coverage, plaque number, and size classes at 100, 150, and 200 days. Sample sizes: *n* = 18 (father) and 20 (mother) at 100 and 200 days; *n* = 20 per group at 150 days. Unpaired two-tailed *t* tests. Means and standard deviations. (B) Representative A*β*-stained sections at 100 days. Scale bar, 1 mm.

**Figure 4:**
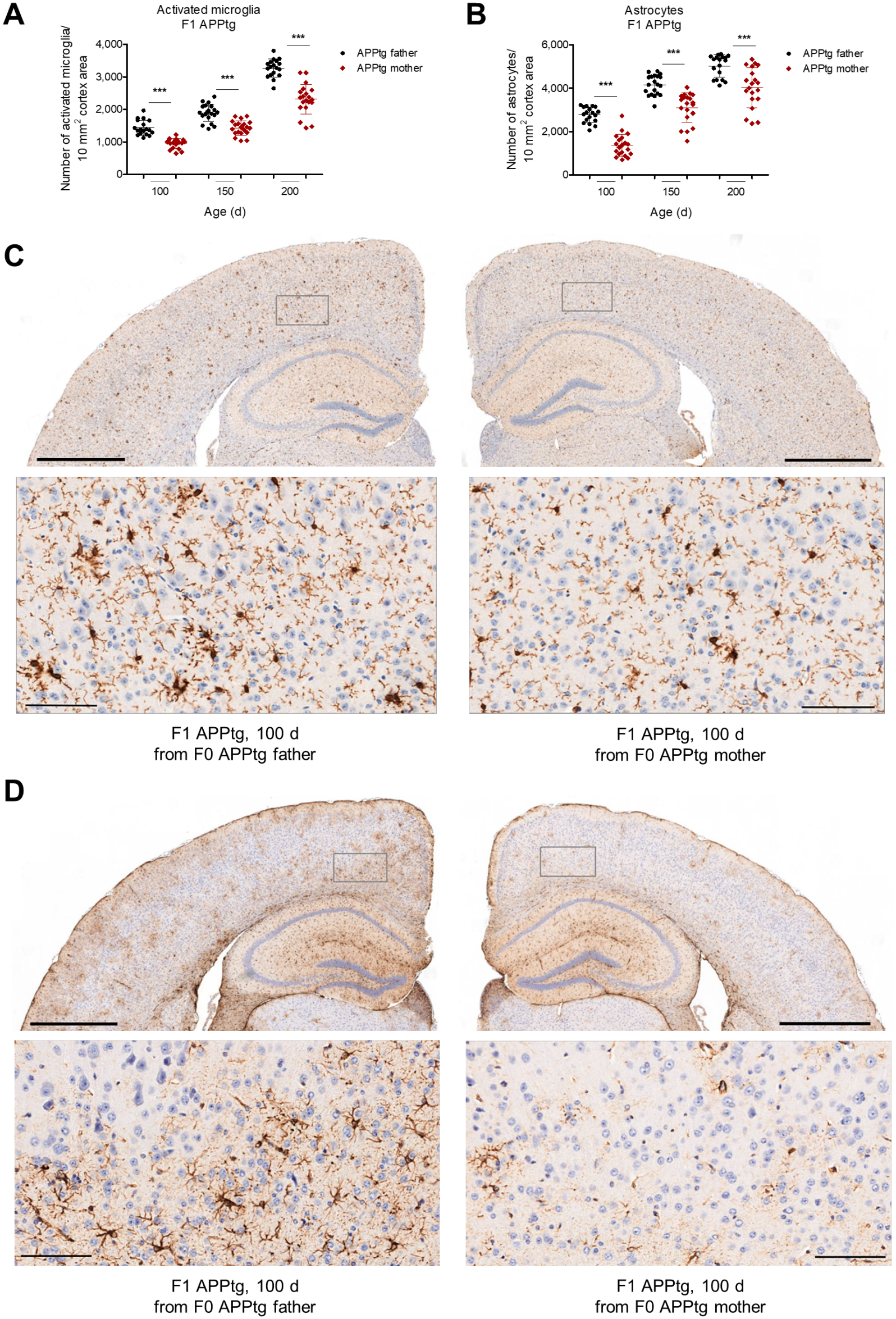
Gliosis is lower after maternal transmission. (A, B) Cortical IBA1-positive microglia and GFAP-positive astrocytes. Sample sizes match Figure 3. Unpaired two-tailed *t* tests. Means and standard deviations. (C, D) Representative sections at 100 days, with insets. Scale bars, 1 mm and 100 μm.

### No A*β*42 difference was detected in the F2 cohort

The comparisons of 100-day-old F2 mice detected no difference in either soluble or in-soluble A*β*42 between the paternal-grandfather and paternal-grandmother classes or between the maternal-grandfather and maternal-grandmother classes (Figure 2). Thus, F2 animals whose father had inherited the transgene from an APPPS1 mother were not distinguishable from those whose father had inherited it from an APPPS1 father. F2 animals born to non-transgenic daughters of APPPS1 mothers also showed no detectable reduction in A*β*42 when they received the transgene from an APPPS1 father. These results provide no evidence that the F1 A*β*42 difference persists through the tested F2 breeding paths.

Three further controls give the same result. First, non-transgenic F1 offspring of APPPS1 mothers and of APPPS1 fathers did not differ in microglia or astrocytes at 100 or 200 days (Figure S8), so APPPS1 mothers do not produce offspring with fewer glia irrespective of the transgene. Second, F0 APPPS1 males and females euthanized at 225 days did not differ in plaques, microglia, or astrocytes (Figure S9), so the F1 difference is not a sex difference already present in the F0 parents. Third, serum antibodies against A*β*42 at 40 days were not higher in offspring of APPPS1 mothers (Figure S10).

### Human APP protein is lower at intermediate ages

Human APP in the membrane fraction was reduced by 62.1% at 100 days (*p <* 0.0001) and by 23.9% at 150 days (*p* = 0.031) after maternal transmission. The same comparison was not significant at 50 or 200 days (*n* = 8 per group; Figure 5). The A*β* difference is therefore present before a detectable APP-protein difference and outlasts it. At the age of peak APP reduction, the protein and insoluble-A*β* changes are of similar size.

**Figure 5:**
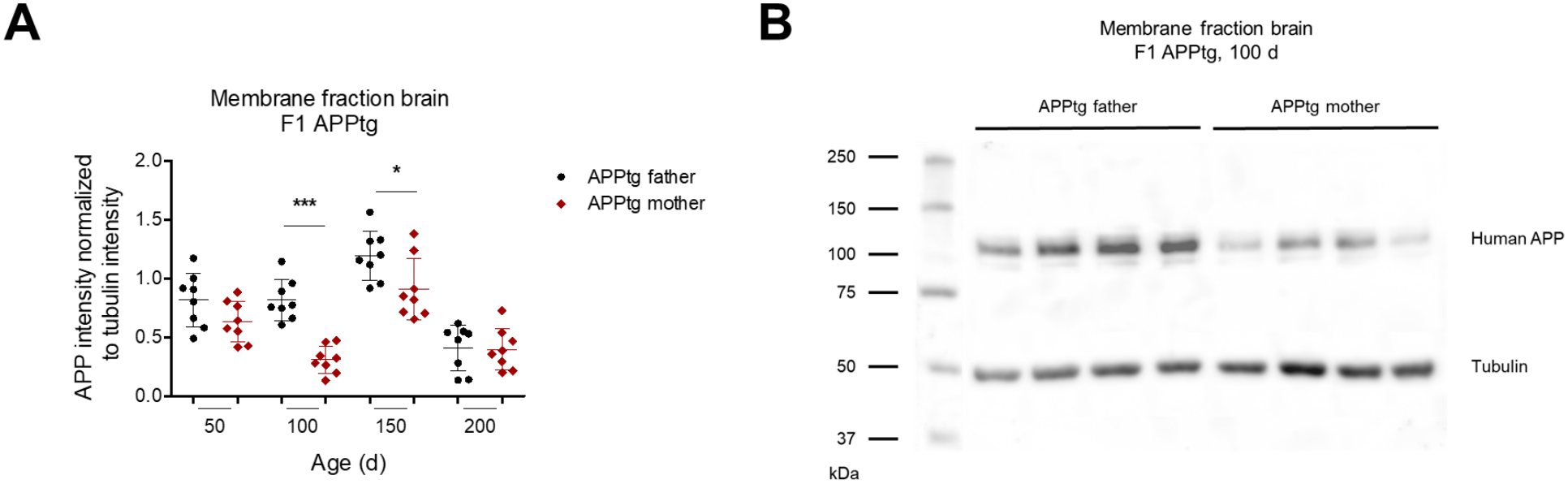
Human APP protein is lower at 100 and 150 days. (A) 6E10 immunoblot of the membrane fraction, normalized to tubulin (*n* = 8 per group and age). Unpaired two-tailed *t* tests. Means and standard deviations. Differences at 50 and 200 days were not significant. (B) Representative blot at 100 days (*n* = 4 per group shown).

### The parent-of-origin methylome is sex-specific and not centered on chromosome 2

The methylation pattern was measured using long-read whole genome sequencing (lr-WGS) in a separate cohort of 32 brains: F0 wild-type and hemizygous males and females, and hemizygous F1 mice with the transgene from the father or the mother (*n* = 4 per sex and parent; Figure S1, Table S1). Differentially methylated positions were called by a permutation false-discovery procedure with an effect-size floor of 20 percentage points.

Pooling F1 males and females identified 6 positions. The same contrast within males returned 66 and within females 37; only four truncated matrix feature identifiers were shared, and the corresponding male and female CpGs were nearby but not always identical (Figure 6, Figure S7). Chromosome X accounted for 32 of 66 male and 19 of 37 female calls (hypergeometric *p* = 1.2 × 10^−21^ and *p* = 5.1 × 10^−14^ against a 6.1% background). Chromosome 2, the insertion chromosome, was not enriched. Three female matrix names labeled chrY:68,889, 68,890, and 68,895 were removed before counting and plotting because those bases are absent from the female bedMethyl files. The eight male chromosome Y calls, at chrY:1,641–64,435, were retained. The first principal component of the matrix separated the sexes. Cross-sex contrasts produced large numbers of sites and were not interpreted as inheritance effects (Figures S2–S6).

**Figure 6:**
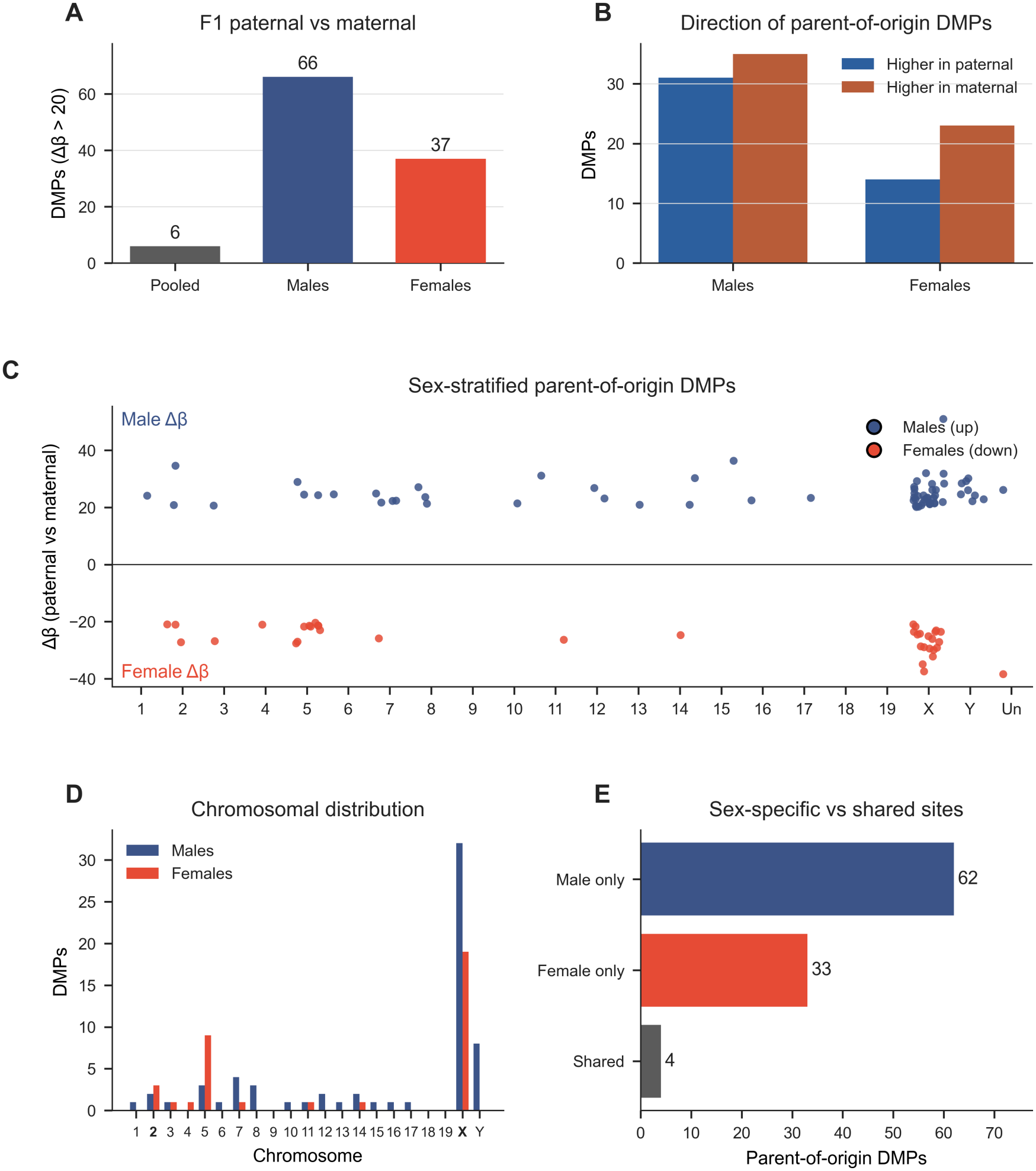
Parent-of-origin methylation in F1 brain is sex-specific. (A) Differentially methylated positions for paternal versus maternal APPPS1 inheritance after permutation-calibrated false-discovery control and an effect-size floor of 20 percentage points. Pooling the sexes yields 6 sites; males yield 66 and females 37. (B) Direction of the sex-stratified differences. (C) Chromosome map, males upward and females downward. (D) Chromo-some X accounts for 32/66 male and 19/37 female calls; chromosome 2 is not enriched. (E) Four truncated matrix feature identifiers are shared between sexes; the mapped CpGs are nearby but not always identical.

Twenty-six of the 103 sex-stratified sites fall in a gene body or promoter (Table S2), including *Pabpc1l2b* and the pseudoautosomal gene *Erdr1x* on chromosome X and the known imprinted genes *Gnas* and *Cdkn1c*. The *Gnas* sites, from chr2:174,140,344 to chr2:174,170,967, are separate from the APPPS1 insertion interval, which maps to 40– 60 cM. Across chromosome 2 in 100 kb bins, the median absolute parent-of-origin difference was about one percentage point, with no shared domain in males and females (Figure S11).

### APP cassette shows a small paternal methylation increase

Methylation of endogenous Thy1 alone would not explain a transgene-specific mechanism. Its promoter (chr9, 43.950–43.955 Mb) differed by less than two percentage points between paternal and maternal F1 inheritance in either sex (Figure S11). *Smo*, which had been noted as a candidate, is on chromosome 6 rather than chromosome 5 and was not a parent-of-origin domain.

The human APP and PSEN1 cDNAs are absent from the mouse reference, so cassette methylation is invisible to a standard bedMethyl contrast. Reads that align to the Thy1 sequence shared with the cassette (promoter through exon 2) and that also contain human APP or PSEN1 sequence report the cassette directly. Every F1 brain yielded such reads (34–63 per mouse). Coverage-weighted 5mC on that shared sequence was higher after paternal transmission for the APP cassette, in both sexes: 17.5% versus 14.4% in males and 17.9% versus 14.5% in females (Figure 7). In each sex the four paternal mice sat above the four maternal mice (Mann–Whitney *p* = 0.029, the smallest *p* this sample size can return). These are nominal comparisons across multiple cassette/sex combinations and should be interpreted as exploratory. The PSEN1 cassette, built on the same Thy1.2 promoter design, did not differ (15.2% versus 13.9% in males, *p* = 0.49; 14.7% versus 14.9% in females, *p* = 0.69). Endogenous reads from the same window also did not differ (*p* = 0.69 in males, *p* = 0.11 in females). The cassette is mostly unmethylated. The small APP-cassette difference does not establish a causal link to the pathology or indicate parent-specific silencing.

**Figure 7:**
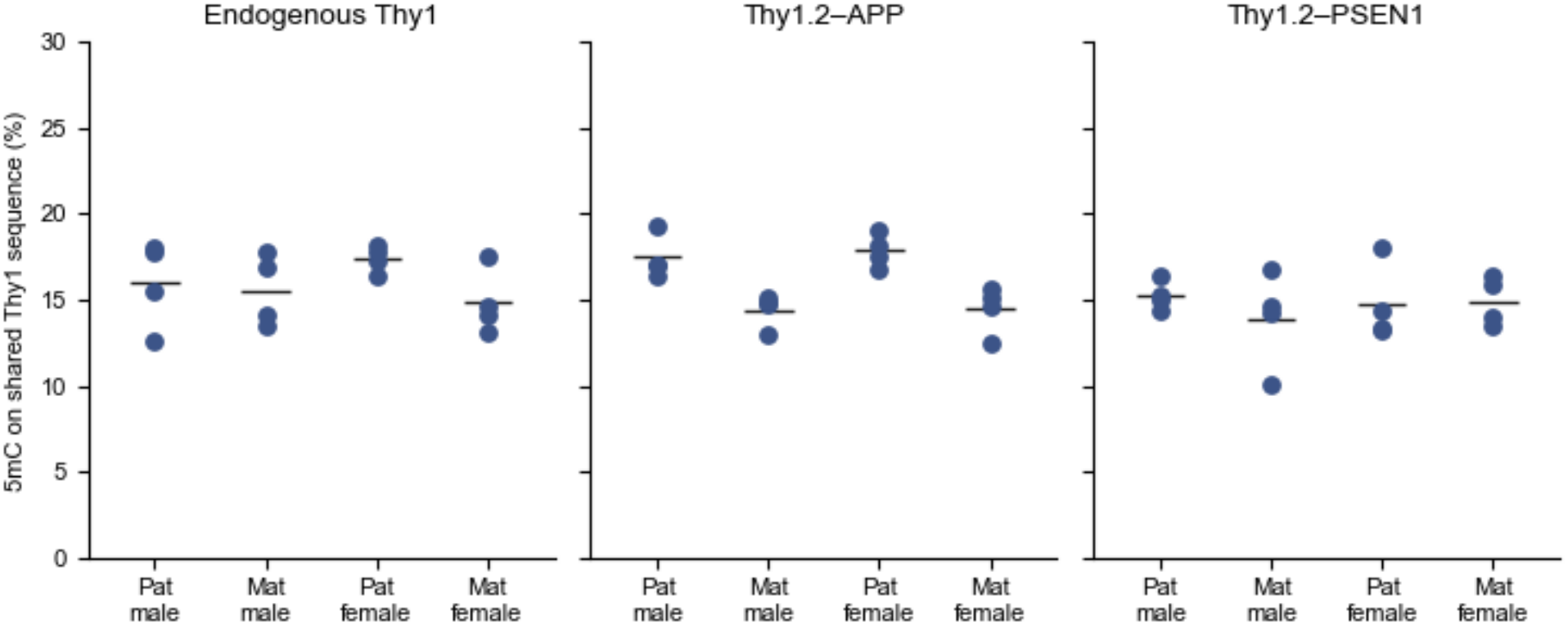
Methylation of the shared Thy1.2 sequence. Each point is one F1 mouse (*n* = 4 per sex and parent). Values are coverage-weighted 5mC on CpGs in chr9:43,950,000 –43,957,315, the promoter-to-exon-2 interval present in both endogenous Thy1 and the Thy1.2 cassette. Cassette reads are those that also contain human APP or PSEN1 sequence. The APP cassette is higher after paternal transmission in both sexes (males 17.5% versus 14.4%, females 17.9% versus 14.5%; Mann–Whitney *p* = 0.029 in each sex). The PSEN1 cassette and endogenous Thy1 reads from the same window do not differ.

## DISCUSSION

These data extend the reported parent-of-origin association in 5xFAD to APPPS1-21, another Thy1-driven APP model.^5,6^ The phenotype here includes soluble and insoluble A*β*42, plaque number and size, microglia, astrocytes, and, at intermediate ages, human APP protein. The A*β*42 difference is present at 50 days and remains detectable at 200 days.

The breeding results narrow, but do not resolve, the mechanism. Lower pathology was observed in transgenic offspring after maternal transmission; the A*β*42 difference was not detected in 100-day-old F2 mice, including offspring of an F1 son that had inherited the transgene maternally. Non-transgenic F1 offspring of APPPS1 mothers did not differ in the measured glial counts, and serum anti-A*β* antibody levels were not elevated at 40 days. Together these observations weigh against some persistent modifiers in the tested crosses but do not rule out mitochondrial, X-linked, maternal-environmental, or other mechanisms. The data are compatible with a parent-of-origin-associated state, but they do not establish whether such a state was transmitted to the tested F2 offspring or reset between generations.

Sasmita et al.^6^ proposed imprinting of the Thy1.2-driven 5xFAD transgene as one possible mechanism. Our data do not show differential methylation of the endogenous Thy1 promoter, and the 100-kb-bin analysis did not identify a broad shared parent-of-origin domain on chromosome 2. The APP cassette had an approximately three-percentage-point paternal methylation increase in both sexes, whereas the co-inherited PSEN1 cassette did not. With four animals per sex and parent-of-origin group, these comparisons are exploratory; they do not establish that the APP-cassette methylation difference changes expression or mediates the pathology. Chromosome X enrichment among the called sites also warrants cautious interpretation given the small methylation cohort and the absence of a detected F2 A*β*42 difference. The practical implication is that APPPS1 studies should record which parent transmitted the transgene and balance or account for breeding direction, which may be a source of variation in plaque burden, gliosis, and APP protein.

## LIMITATIONS OF THE STUDY

Phenotype comparisons pool males and females and test each age separately. The phenotype *p* values are unadjusted for testing across ages and outcomes and should be interpreted as nominal. Litter was recorded but was not a random effect in the models used here. The methylation cohort comprises 32 brains, with four F1 mice per sex and parent-of-origin group. Of the 103 sex-stratified calls, 26 fall in a gene body or promoter (Table S2) and 50 lie in the first 100 kb of a chromosome, before the first annotated GRCm39 gene. The other 27 were not placed, because no CpG in the truncated window matched the matrix means within 15 percentage points in at least three mice of each parental arm. Twenty-three of the 26 placed sites met that match only after the coverage requirement was lowered from five reads to one, so those gene assignments are less certain than the three sites covered by at least five reads. The APPPS1 insertion is known only as a genetic interval on chromosome 2, not as a base-pair coordinate, so flanking sequence of the integration site was not tested directly. Cassette methylation was called from junction-spanning nanopore reads rather than from an assembled transgene contig, and it was not confirmed by bisulfite sequencing. The F2 cohort was assessed for A*β*42, not histology or APP protein.

## RESOURCE AVAILABILITY

### Lead contact

Requests for further information and resources should be directed to Jens Pahnke.

### Materials availability

This study did not generate new unique reagents. APPPS1-21 mice were obtained from the University of Tübingen. Fixed tissue, frozen hemispheres, and protein extracts from the cohorts described here are archived in the Pahnke laboratory.

### Data and code availability

- Nanopore base calls, bedMethyl files, and the methylation analysis code used for Figures 6–7 and S1–S11 are archived on ZENODO (10.5281/zenodo.24392751).
- Any additional information required to reanalyze the data will be available from the lead contact upon request.

## ACKNOWLEDGMENTS

The authors thank Prof. Einar Vik-Mo from the Department of Neurosurgery Research (OUS) for access to Nanopore sequencing instrumentation.

## AUTHOR CONTRIBUTIONS

Conceptualization, M.B. and J.P.; data curation, M.B., S.H., J.G., and J.P.; formal analysis, M.B., S.H., J.G., and J.P.; funding acquisition, J.P.; investigation, M.B. and J.P.; methodology, M.B., T.B., and I.E.; project administration, J.P.; resources, J.P.; software, J.G. and J.P.; supervision, J.P.; validation, M.B. and J.P.; visualization, M.B., J.G., and J.P.; writing – original draft, M.B. and J.P.; writing – review and editing, all

## DECLARATION OF INTERESTS

The authors declare no competing interests.

### DECLARATION OF GENERATIVE AI AND AI-ASSISTED TECHNOLOGIES

During the preparation of this work, the authors used Grok (different models) in Cursor (SpaceXAI) and ChatGPT (different models) in PrismAI (OpenAI) to generate analysis code, organize source materials and to assist with text editing and L^A^TEX. After using these tools, the authors reviewed and edited the content as needed and take full responsibility for the content of the publication.

## STAR⋆METHODS

### Key resources table

| REAGENT or RESOURCE | SOURCE | IDENTIFIER |
|---|---|---|
| **Antibodies** |  |  |
| Anti-β-amyloid, 17–24, clone 4G8 | BioLegend | Cat# 800703 |
| Anti-β-amyloid, 1–16, clone 6E10 | BioLegend | Cat# 803003 |
| Anti-Iba1 | FUJIFILM Wako | Cat# 019-19741 |
| Anti-GFAP | Agilent | Cat# Z033401-2 |
| Anti-β-tubulin isotype III, clone SDL.3D10 | Merck | Cat# T5076 |
| HRP goat anti-mouse IgG | Bethyl Laboratories | Cat# A90-216P |
| Anti-mouse antibody (goat), SULFO-TAG labeled | Meso Scale Diagnostics | Cat# R32AC-5 |
| **Chemicals, peptides, and recombinant proteins** |  |  |
| Aβ42, ultrapure, NH<sub>4</sub>OH | rPeptide | Cat# A-1167 |
| RNAlater | Merck | Cat# R0901-500ML |
| IGEPAL | Merck | Cat# 9002-93-1 |
| cOmplete, Mini Protease Inhibitor Cocktail | Roche | Cat# 4693124001 |
| Pierce BCA Protein Assay Kit | Thermo Fisher Scientific | Cat# 23225 |
| 4× Laemmli sample buffer | Bio-Rad | Cat# 1610747 |
| **Critical commercial assays** |  |  |
| V-PLEX Plus Aβ42 Peptide (4G8) Kit | Meso Scale Diagnostics | Cat# K150SLE-4 |
| DNeasy Blood & Tissue Kit | Qiagen | Cat# 69504 |
| Qubit dsDNA HS Assay Kit | Thermo Fisher Scientific | Cat# Q32851 |
| PromethION R10.4.1 flow cell | Oxford Nanopore Technologies | RRID:SCR_017987 |
| **Experimental models: Organisms/strains** |  |  |
| APPPS1-21 | University of Tübingen | MGI:3765351 |
| C57BL/6J | JAX Labs | RRID:IMSR_JAX:000664 |
| GRCm39 | NCBI RefSeq | GCF_000001635.27 |
| **Deposited data** |  |  |
| Nanopore base calls, alignments, and bedMethyl, Methylation and cassette-calling code | This paper | 10.5281/zenodo.24392751 |
| **Software and algorithms** |  |  |
| Prism 6 | GraphPad | RRID:SCR_002798 |
| DeePathology STUDIO | DeePathology.AI | N/A |
| ImageJ | NIH | SCR_003070 |
| MinKNOW | Oxford Nanopore Technologies | RRID:SCR_023196 |
| Dorado | Oxford Nanopore Technologies | RRID:SCR_025883 |
| minimap2 | Heng Li | RRID:SCR_018550 |
| modkit | Oxford Nanopore Technologies | RRID:SCR_028163 |
| **Devices** |  |  |
| MESO QuickPlex SQ120 | Meso Scale Diagnostics | RRID:SCR_020304 |
| PromethION 24 | Oxford Nanopore Technologies | RRID:SCR_017987 |
| BOND-III stainer | Leica Biosystems | RRID:SCR_026521 |
| Pannoramic Midi II | 3DHistech | RRID:SCR_024834 |
| Octoplus QPLEX | DyeAGNOSTICS | N/A |

### Experimental model and study participant details

#### Mice

APPPS1-21 mice co-express human APP (KM670/671NL) and PSEN1 (L166P), each driven by mouse Thy1.2. The line was obtained from the University of Tübingen and maintained hemizygous on C57BL/6J.^4^ The allele is listed as MGI:3765351. Non-transgenic partners were C57BL/6J mice from the same colony. Experiments followed European Directive 2010/63/EU and Norwegian law and were approved by the national animal-research authority (Mattilsynet FOTS 7049, 7771, and 20540).

Animals were bred and housed at the Department of Comparative Medicine (Radium Hospital section), Oslo University Hospital, Norway. Macroenvironmental conditions were an average temperature of 22 ± 1^°^C, relative humidity of 62 ± 5%, ventilation of 15 air changes per hour, and a 12 h dark/12 h light cycle. Illuminance was 1 lux at night, 70 lux during the day, and 400 lux for working illumination. Mice were kept in Eurostandard type III open Makrolon® cages with aspen bedding (*Populus tremula*; Tapvei®, Estonia) and enrichment (tissue paper, tunnels, rods, and occasionally gnawing sticks). Cages were changed weekly and housed up to eight mice. Food was provided as expanded maintenance pellets (SDS, Estonia), available ad libitum, and drinking water was acidified to pH 3. Health monitoring was performed at least three times a year: one annual screen covered the full FELASA-recommended panel, and the other two targeted selected microorganisms. Until these experiments, the colony was maintained under SOPF (specific opportunistic pathogen-free) conditions. Segmented filamentous bacteria were the only microorganism detected.

Both males and females were studied. Ages were 225 days for the F0 histology cohort, 40, 50, 100, 150, or 200 days for F1 mice, and 100 days for F2 mice. Phenotype comparisons pool the sexes within each parent-of-origin group. A sex-by-parent interaction was not tested, so the amyloid results do not establish that the parent-of-origin effect is the same size in males and females. The one direct sex comparison is F0 APPPS1 males versus females at 225 days (Figure S9). Methylation was analyzed within sex (Table S1). No a priori sample-size calculation is recorded. Mice were included by genotype and by which parent transmitted the transgene. No exclusions are recorded. Groups were the breeding classes in Figure 1, not randomized treatment arms. Subsets used for A*β*42, the APP immunoblot, and non-transgenic histology were drawn at random from those classes. Sample sizes are the *n* values in the figure legends and in Table S1.

## Method details

### Breeding

F0 APPPS1 males were crossed to non-transgenic C57BL/6J females, and F0 non-transgenic males were crossed to F0 APPPS1 females (Figure 1). Every F0 APPPS1 breeder had inherited the transgene from an APPPS1 father. For the F2 generation, F1 APPPS1 males of either parental origin were crossed to non-transgenic females, and F1 non-transgenic females of either parental origin were crossed to APPPS1 males that had inherited the transgene from their father.

### Tissue collection

Mice received a ketamine/xylazine overdose (400 mg/kg and 40 mg/kg) and were perfused intracardially with 20 ml of cold PBS. One hemisphere was snap-frozen. The other was fixed in 4% paraformaldehyde. Blood from the caudal vena cava was allowed to clot at room temperature for at least 1 h, centrifuged at 2,000 *× g* for 15 min at 4^°^C, and the serum stored at −80^°^C.

### A*β*42 immunoassay

Frozen hemispheres from F1 and F2 APPPS1 mice were thawed on ice in RNAlater for 1 h and homogenized for 60 s.^7^ Soluble and insoluble A*β* were extracted in TBS (10 μl per mg homogenate) and then in 5 M guanidine (8 μl per mg). A*β*42 was quantified with the V-PLEX Plus A*β*42 Peptide (4G8) kit on a MESO QuickPlex SQ120 according to the manufacturer’s instructions and normalized to sample weight. The experimenter was blinded to group.

### Immunohistochemistry

Fixed hemispheres were paraffin-embedded and cut at 4 μm. Sections at approximately 0.14, −1.7, and − 2.5 mm from bregma were stained on a BOND-III system for A*β* (4G8, 1:4,000), IBA1 (1:1,000), and GFAP (1:500).^7,8^ Slides were scanned at 230 nm resolution on a Pannoramic Midi II. DeePathology STUDIO was trained separately to recognize plaques, activated microglia, and astrocytes.^9,10^ After that training, the cortex (parietal, somatosensory, auditory, and temporal) was outlined and the sections were scored automatically. Counts were normalized to 10 mm^2^. The experimenter was blinded to group.

### Human APP immunoblot

About 30 mg of homogenate from F1 APPPS1 mice was extracted in two detergent steps.^11^ The first buffer was 200 μl of 50 mM Tris-HCl, pH 7.4, 150 mM NaCl, 0.1% Triton X-100, and protease inhibitor. After homogenization for 30 s and centrifugation at 14,000 × *g* for 90 min at 4^°^C, the pellet was extracted in 100 μl of 50 mM Tris-HCl, pH 7.4, 137.5 mM NaCl, 5 mM EDTA, 1 mM PMSF, 0.5% Triton X-100, and 1% IGEPAL, and centrifuged under the same conditions. Protein in the second supernatant was measured with a Pierce BCA assay. Sixty micrograms of protein were mixed with 4 × Laemmli buffer, separated on a 7.5% gel (15 min at 100 V, then 40 min at 200 V), and transferred to 0.45 μm PVDF. Membranes were blocked for 1 h at room temperature in 1.5% non-fat dry milk in PBS with 0.01% Tween-20, probed with 6E10 (1:1,000) and anti-*β*-tubulin iso-type III (1:5,000), and detected with HRP goat anti-mouse IgG (1:5,000) on an Octoplus QPLEX imager. Band intensity was measured in ImageJ and APP was normalized to tubulin. The experimenter was blinded to group.

### Serum anti-A*β* assay

Ultrapure A*β*42 was resuspended in 1% NH_4_OH at 1 mg/ml and sonicated on ice twice for 30 s. QuickPlex 96-well plates were spot-coated overnight at room temperature with 10 μg/ml A*β*42 in PBS containing 0.03% Triton X-100, then blocked for 1 h with MSD Blocker A. Serum was diluted fourfold in Diluent 100. A 4G8 calibration curve was prepared in the same diluent, from 400 ng/μl to 0.097 ng/μl, with diluent as the zero standard. After washing in PBS with 0.05% Tween-20, calibrators and samples were incubated for 2 h at room temperature. Bound mouse immunoglobulin was detected with SULFO-TAG anti-mouse antibody (1:500, 1 h), and the plates were read in 2 × Read Buffer T on a MESO QuickPlex SQ120. The experimenter was blinded to group.

### DNA extraction

DNA was extracted from frozen brain tissue of the 32 mice in Table S1 with the DNeasy Blood & Tissue Kit (Qiagen) according to the manufacturer’s tissue protocol. DNA was quantified with the Qubit dsDNA HS assay (Thermo Fisher Scientific) and normalized to 100 ng/μl.

### Nanopore methylation

Thirty-two brains (Table S1) were sequenced on PromethION R10.4.1 flow cells on a PromethION 24 instrument running MinKNOW. Raw reads were base-called with Dorado (v.7.3.9) in the high-accuracy setting, and 5mC was called in the CG context. Base-called reads were aligned to GRCm39 (GCF_000001635.27) with minimap2 (v.2.25). Modified-base calls were summarized as bedMethyl files with modkit (v.0.2.7). The working feature matrix contained 2,619,370 positions. Percentages were converted to fractions (*β* = %*/*100). For each position the effect size was the absolute difference in group means. Group labels were permuted 1,000 times with group sizes held constant. At each candidate threshold, the estimated false-discovery rate was the mean number of permuted positions passing that threshold, divided by the number of observed positions passing it. A position was retained at the smallest threshold whose estimated false-discovery rate was at most 0.05, and only if the absolute difference also exceeded 20 percentage points. No variance or missingness filter was applied before testing. After the calls, three female parent-of-origin features (NC_000087.868889, NC_000087.868890, and NC_000087.868895) were removed because those bases are absent from the female bedMethyl files; the matrix had stored the missing positions as zero. Sex-stratified and pooled contrasts were run separately. Principal-component analysis used the 80,000 most variable features.

Matrix feature names are limited to 17 characters, which cuts coordinates of seven or more digits. For a cut-off name, bedMethyl CpGs in the window still consistent with the retained digits were compared with the matrix means. The CpG with the smallest combined absolute error was kept when that error was at most 15 percentage points and at least three mice contributed to each parental arm. The comparison required coverage of at least five reads and was repeated at one read when the five-read comparison did not meet the error limit. Names shorter than 17 characters were used as stored. Gene overlap was read from the GRCm39 RefSeq annotation. A site was classified as promoter-associated if it fell within 2 kb upstream of a transcription start site and outside all annotated gene bodies. Coordinate-level effect sizes can differ from the matrix-feature values used for the 20-percentage-point selection floor. Endogenous *Thy1, Smo*, and 100 kb bins of chromo-somes 2 and 5 were summarized directly from bedMethyl as coverage-weighted percent methylation.

### Cassette methylation

The Thy1.2 cassette is not in the reference. Primary alignments to chr9:43,940,000– 43,970,000 were inspected. A read was assigned to the APP or PSEN1 cassette when its sequence contained at least two non-overlapping 15-mers from the corresponding human coding sequence. 5mC was taken from the MM/ML tags (C+m, probability at least 0.5) on CpGs in the shared Thy1 interval chr9:43,950,000–43,957,315. For reverse-strand alignments the MM tag follows the sequenced orientation rather than the stored reverse complement; those positions were converted before counting. Mouse-level values are coverage-weighted percentages.

### Quantification and statistical analysis

#### Phenotype statistics

Phenotype tests were run in Prism 6. The exact *n*, the measure of center, and the measure of dispersion are given in the figure legends. *n* is the number of mice. Normality was assessed with the D’Agostino–Pearson test, and a normal distribution was rejected at *p <* 0.05. F1 A*β*42 comparisons, F0 male–female histology, and non-transgenic microglia comparisons used two-tailed Mann–Whitney tests. F1 histology, F1 APP immunoblots, F2 A*β*42, and non-transgenic astrocyte comparisons used unpaired two-tailed *t* tests. Serum antibody concentrations were compared by a Kruskal–Wallis test. Tests were performed within each age, with no adjustment across ages or outcomes. *p <* 0.05 was treated as significant; phenotype *p* values should be interpreted as nominal. No litter term was fitted, and sexes were pooled.

#### Methylation calls

The permutation rule, the 20-percentage-point floor, and the coordinate-rescue rule are described under Method details. Chromosome enrichment compared the share of called positions on a chromosome with the share of matrix features on that chromosome, using a hypergeometric test. Cassette methylation was compared within sex by a two-sided

Mann–Whitney test. Exact *n*, and what *n* represents, is given in Figure 7 and Table S1.

## SUPPLEMENTAL FIGURES

**Figure S1:**
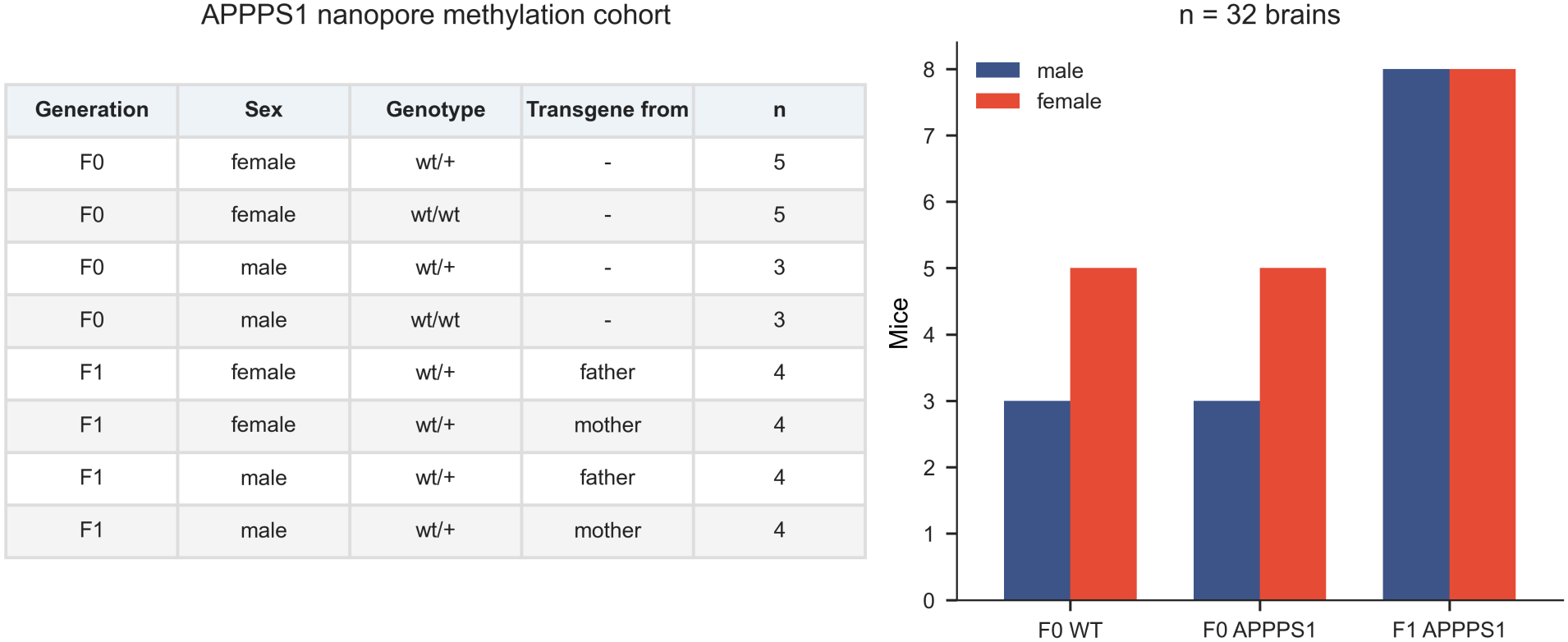
Nanopore methylation cohort, related to Figure 6. Thirty-two brains. F0 wild-type and APPPS1 mice are 3 males and 5 females in each genotype class. F1 mice are hemizygous, with four males and four females for each parent of origin. There are no F1 wild-type animals in the methylation cohort.

**Figure S2:**
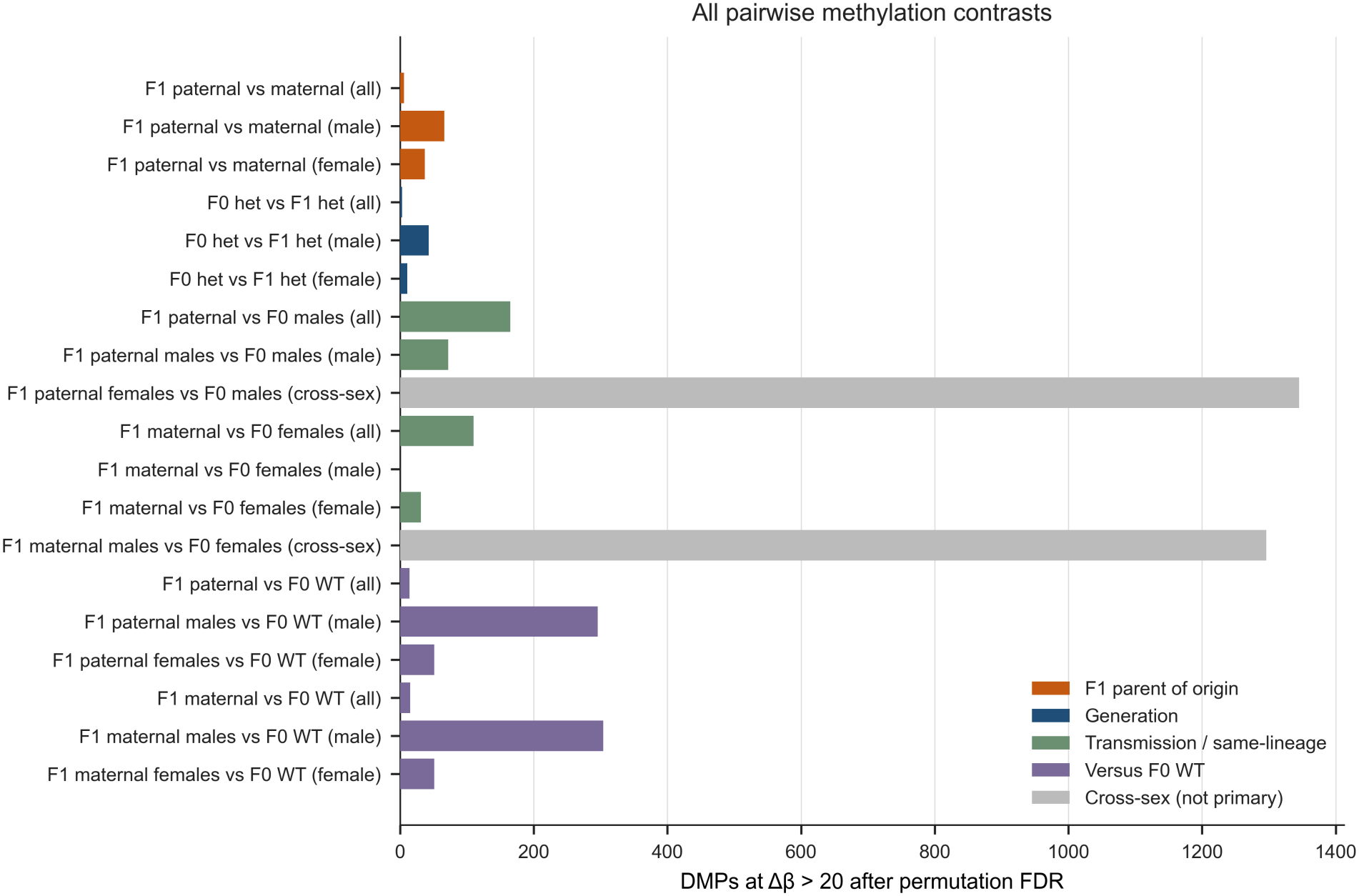
All pairwise methylation contrasts, related to Figure 6. Counts of positions passing the same permutation threshold. Grey bars are cross-sex or otherwise confounded contrasts and are not interpreted as parent-of-origin effects.

**Figure S3:**
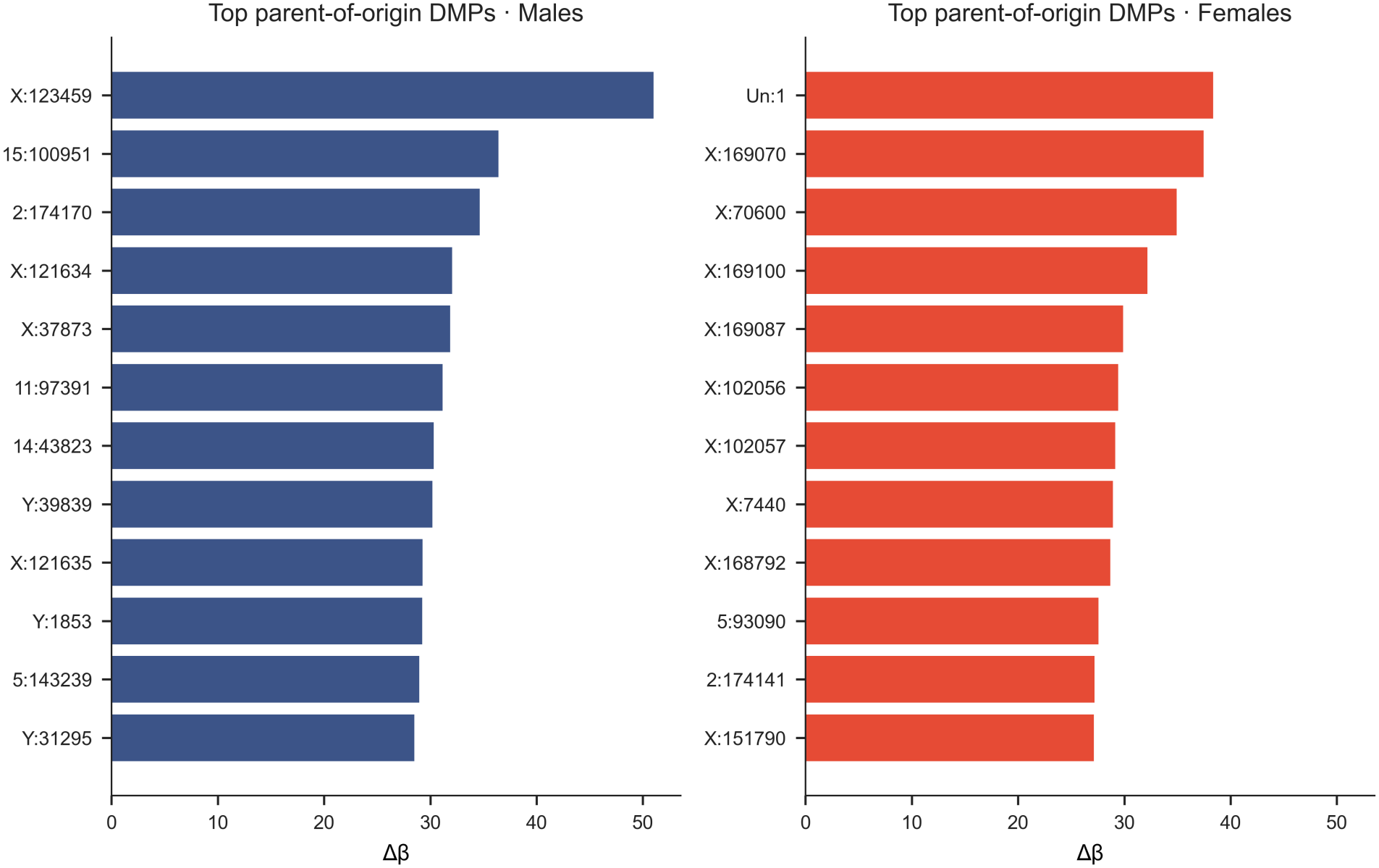
Largest sex-stratified parent-of-origin differences, related to Figure 6. Top positions in F1 males and females, ranked by the paternal-minus-maternal difference. Feature names are the truncated matrix identifiers.

**Figure S4:**
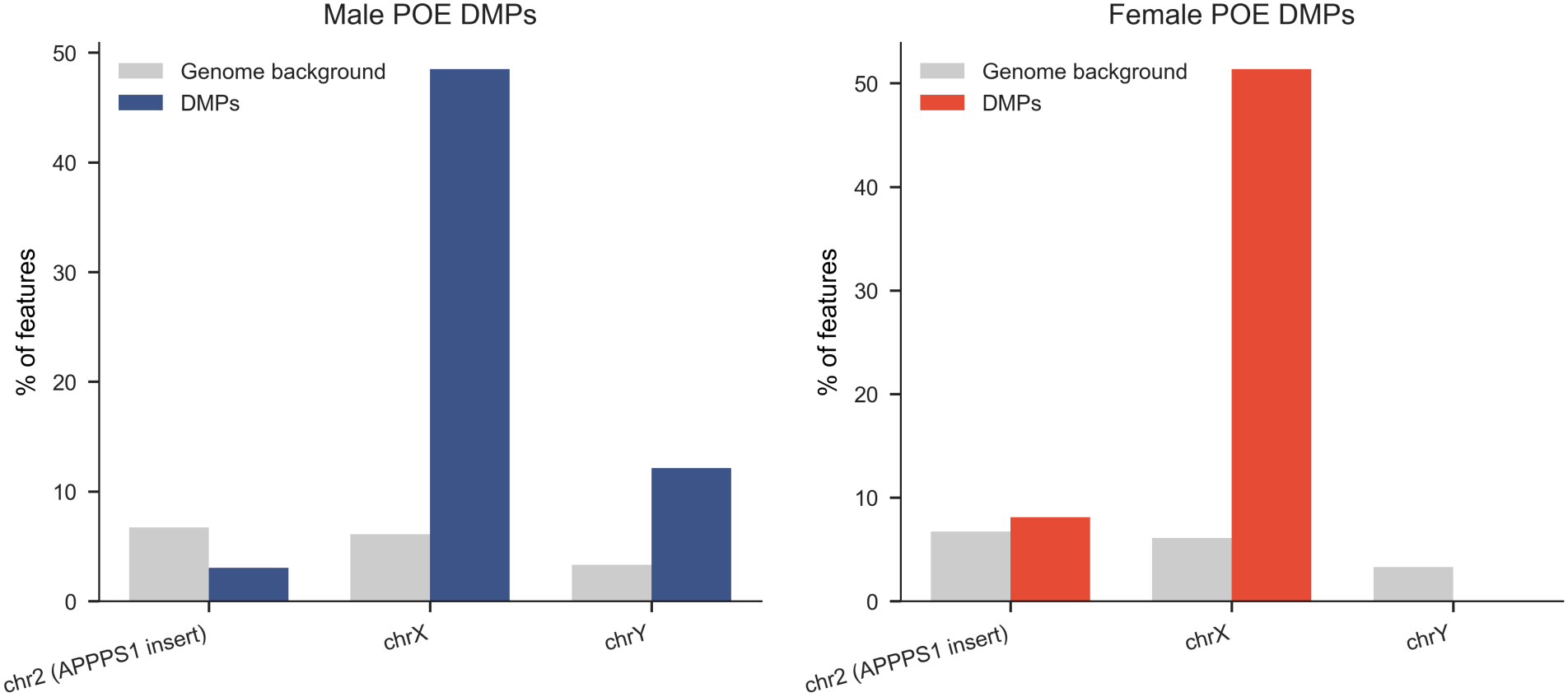
Chromosome 2 is not enriched; chromosome X is, related to Figure 6. Observed share of parent-of-origin calls versus the genomic background of the feature matrix. Chromosome 2 carries the APPPS1 insertion.

**Figure S5:**
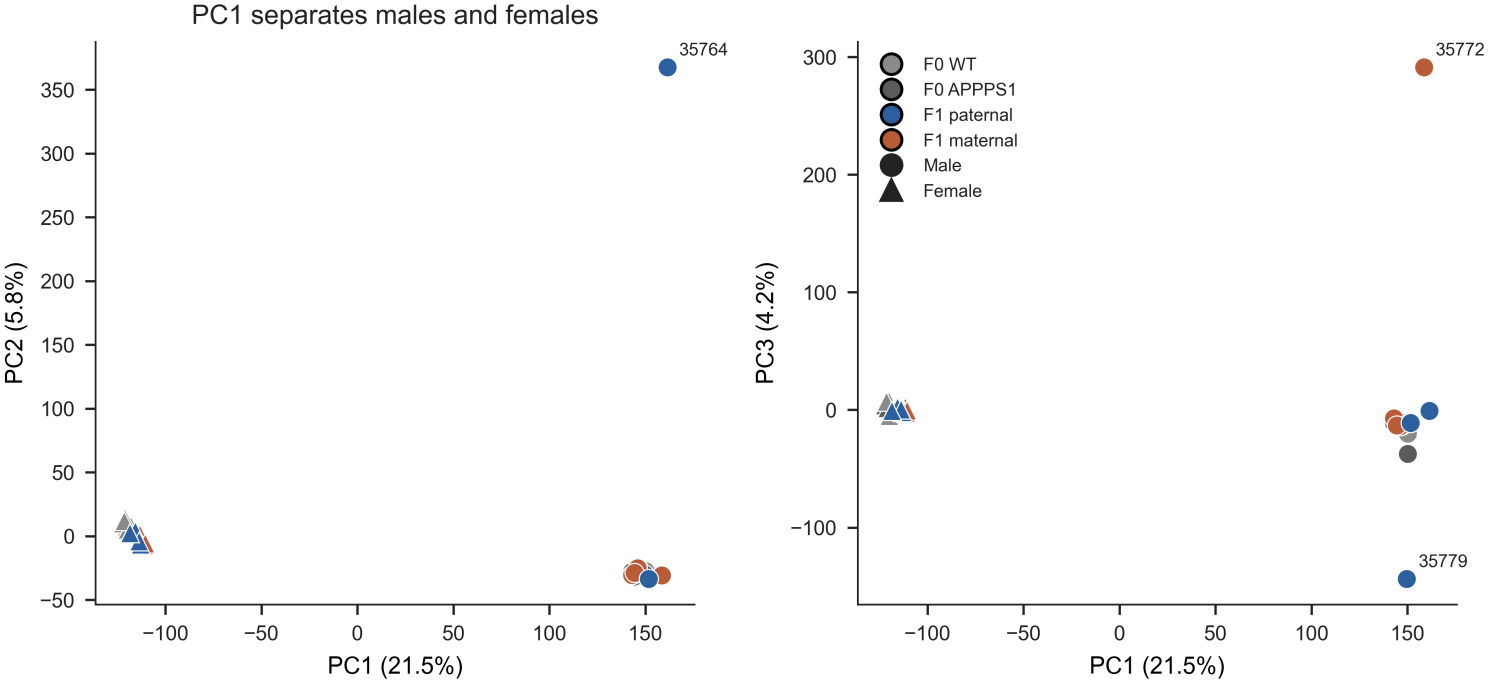
The methylation matrix separates samples by sex, related to Figure 6. Principal components of 32 samples on the 80,000 most variable features. PC1 is sex. Circles, males; triangles, females. Three F1 males (35764, 35772, and 35779) are outliers on later components.

**Figure S6:**
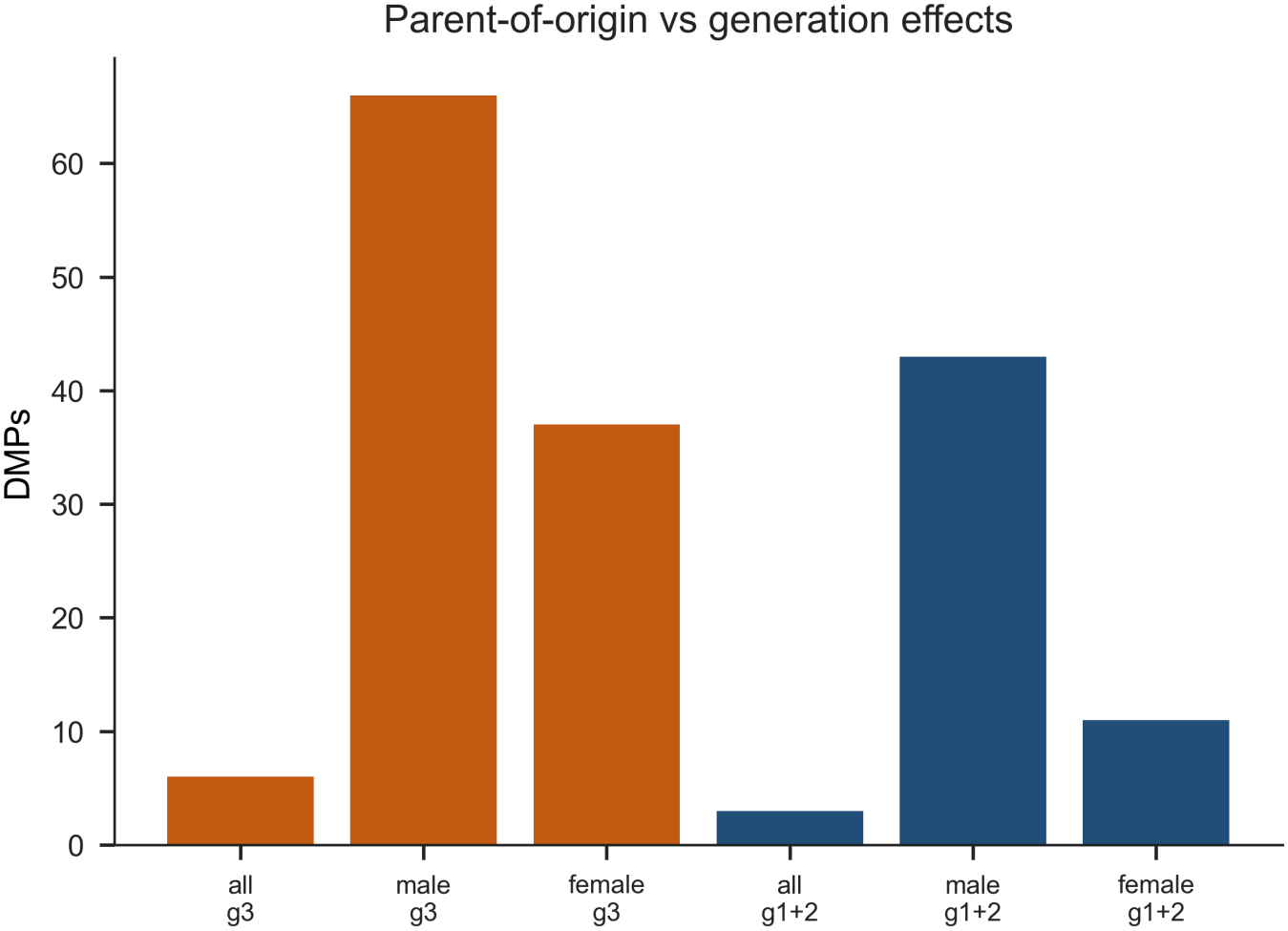
Parent-of-origin and generation contrasts yield different call counts, related to Figure 6. Counts for F1 paternal versus maternal inheritance compared with F0 versus F1 contrasts.

**Figure S7:**
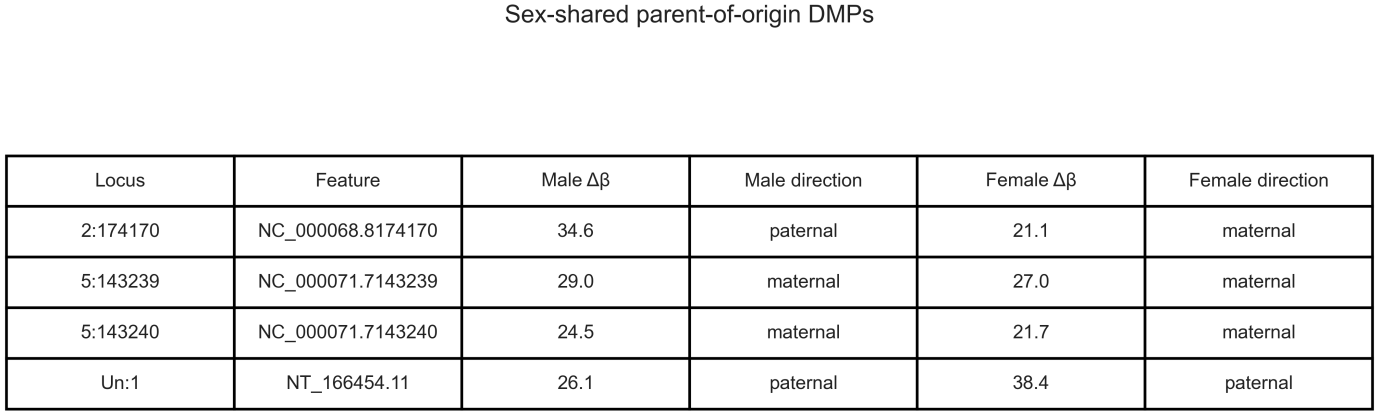
The four shared parent-of-origin feature identifiers, related to Figure 6. The chromosome 2 feature maps to *Gnas* in both sexes, and both chromosome 5 features map to *Zfp316*. Within each identifier the male and female CpGs are nearby rather than identical. The fourth identifier lies on an unplaced contig and has no gene assignment.

**Figure S8:**
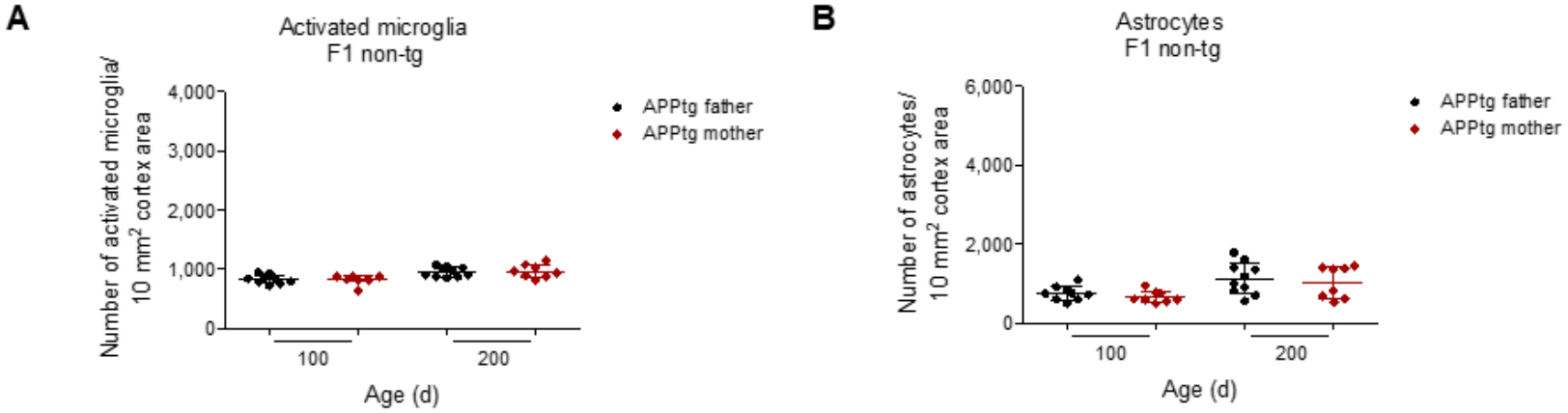
Non-transgenic offspring do not differ by the APP status of the mother, related to Figure 4. (A) Cortical IBA1 microglia and (B) GFAP astrocytes in F1 non-transgenic mice from APPPS1 fathers or mothers. Father group, *n* = 9 at 100 days and *n* = 10 at 200 days; mother group, *n* = 8 at both ages.

**Figure S9:**
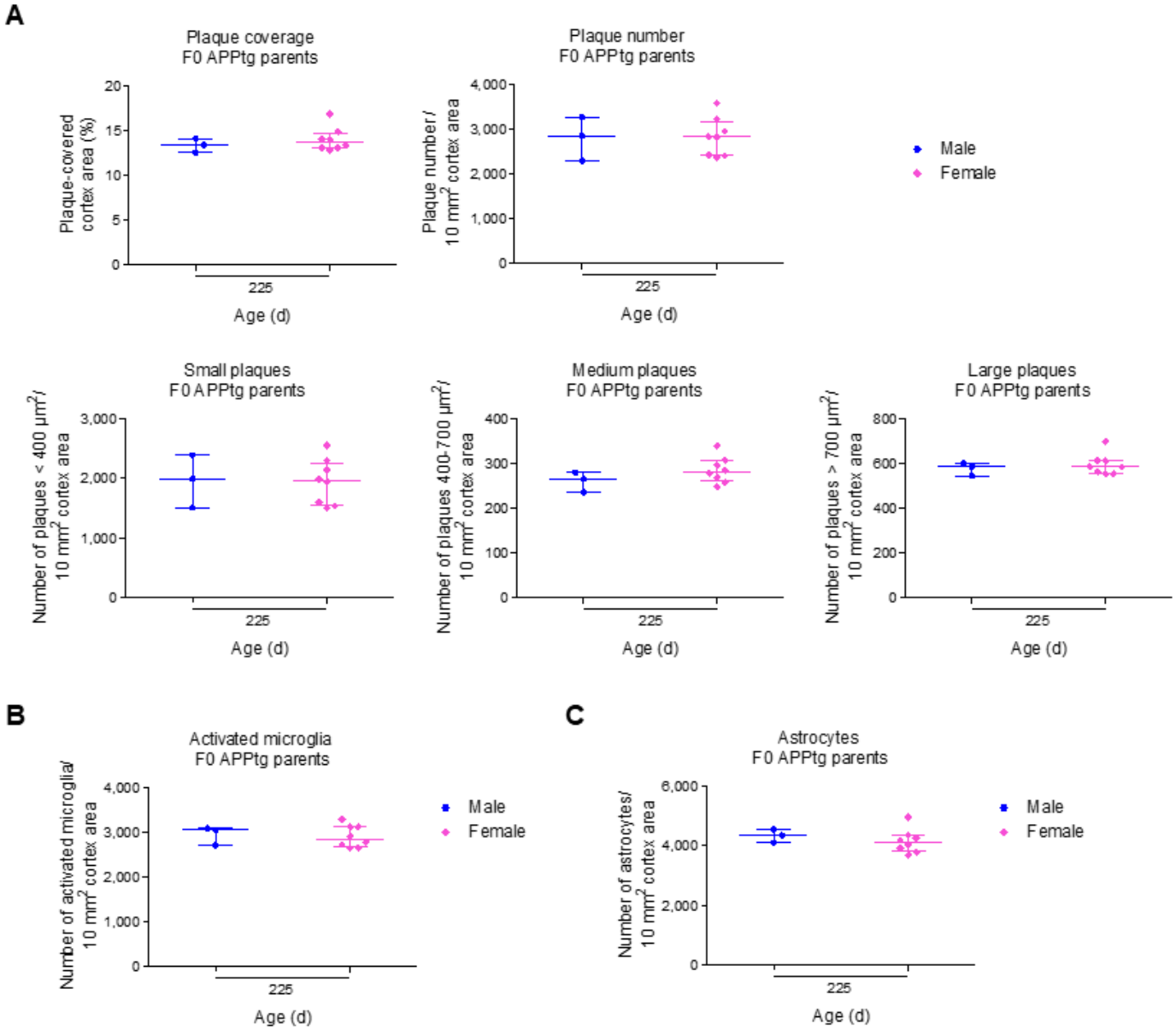
F0 APPPS1 males and females do not differ in plaques or gliosis, related to Figures 3 and 4. (A) Cortical 4G8 plaque measures, (B) IBA1 microglia, and (C) GFAP astrocytes at 225 days (*n* = 3 males, *n* = 8 females). Medians and interquartile ranges.

**Figure S10:**
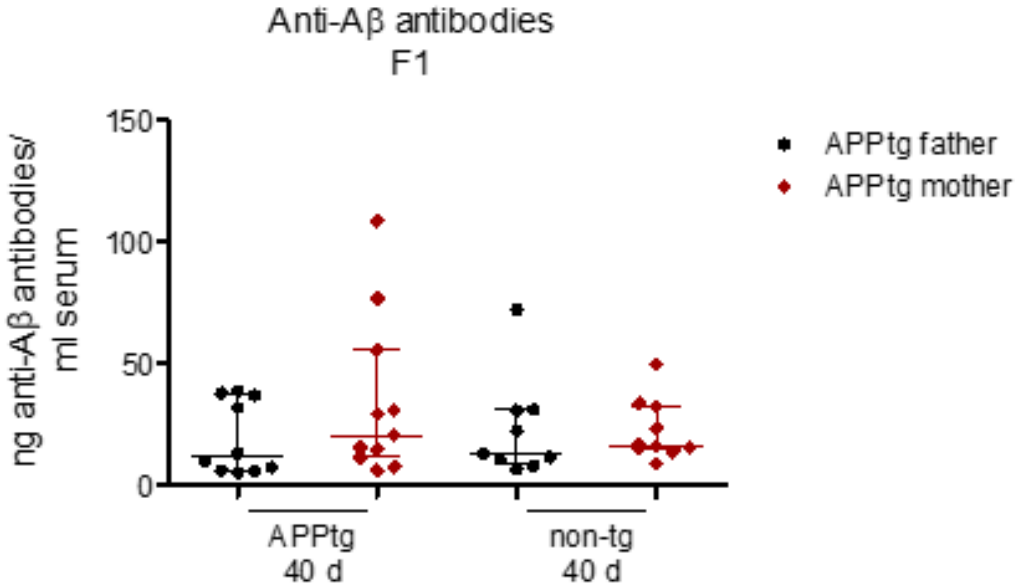
Serum antibodies against A*β* are not higher after maternal transmission, related to Figure 2. Anti-A*β* immunoglobulin in serum at 40 days. Groups are APPPS1 and non-transgenic offspring of APPPS1 fathers or mothers (*n* = 9–11). Medians and interquartile ranges. Kruskal–Wallis test.

**Figure S11:**
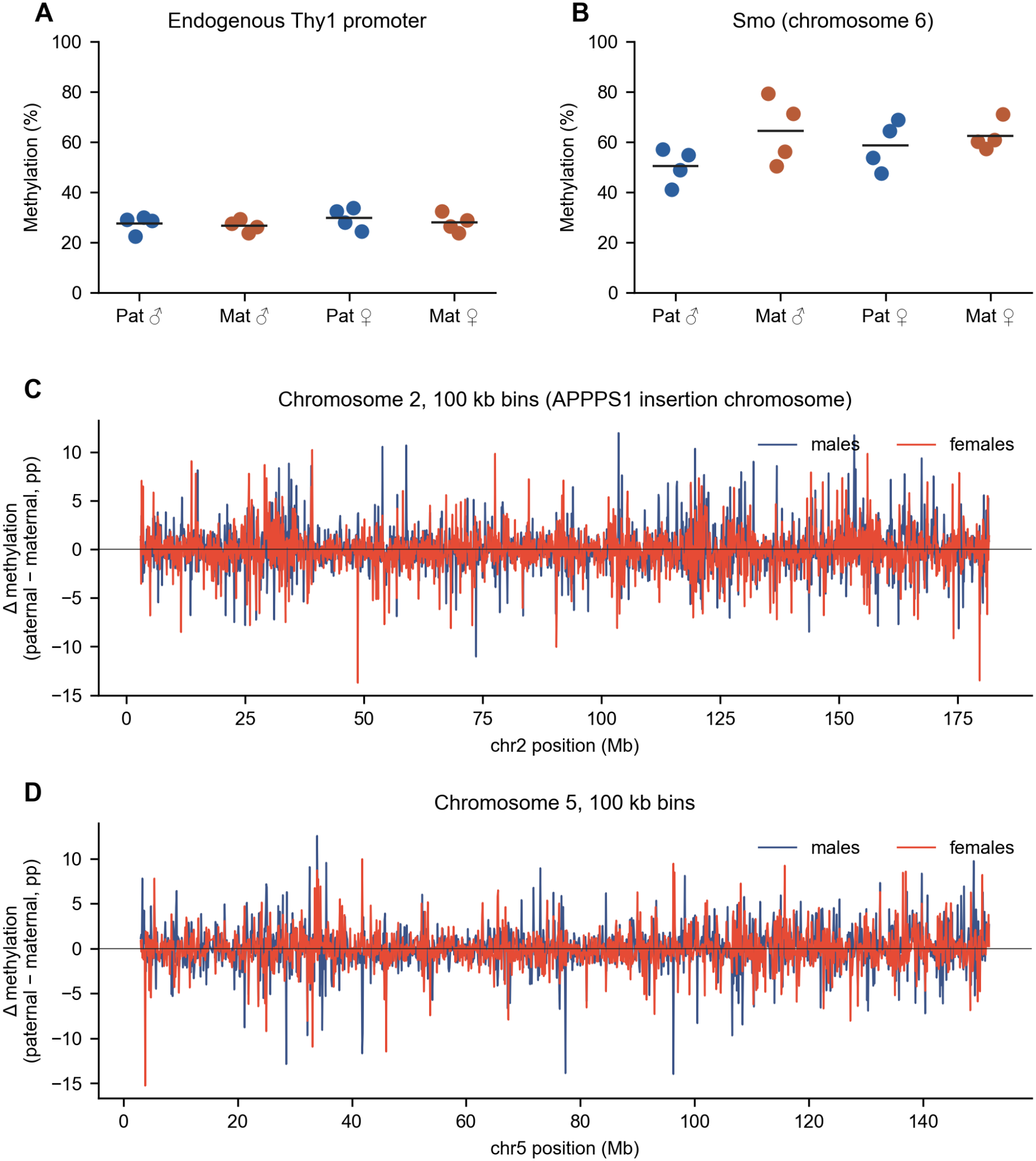
Endogenous Thy1, *Smo*, and chromosomes 2 and 5, related to Figure 7. **(A)** Endogenous Thy1 promoter, coverage-weighted methylation from bedMethyl. Paternal versus maternal F1 inheritance differs by less than two percentage points in either sex. (B) *Smo*, on chromosome 6. (C, D) Paternal-minus-maternal difference in 100 kb bins on chromosomes 2 and 5. The median absolute difference is about one percentage point.

## SUPPLEMENTAL TABLES

**Table S1:** Nanopore methylation cohort, related to Figure 6. F0 animals are the parental generation. F1 animals are hemizygous offspring. Wild-type F1 mice were not sequenced.

| Generation | Sex | Genotype and transgene parent | *n* |
|---|---|---|---|
| F0 | male | APPPS1, paternal | 3 |
| F0 | female | APPPS1, paternal | 5 |
| F0 | male | wild type | 3 |
| F0 | female | wild type | 5 |
| F1 | male | APPPS1 from father | 4 |
| F1 | female | APPPS1 from father | 4 |
| F1 | male | APPPS1 from mother | 4 |
| F1 | female | APPPS1 from mother | 4 |

**Table S2:** Sex-stratified parent-of-origin positions placed on a gene or promoter, related to Figure 6. A truncated matrix feature was matched to the bedMethyl CpG whose paternal and maternal means were closest to the matrix means, with at least three mice per arm. The search used CpGs covered by at least five reads and was repeated at one read when the five-read search missed the matrix means by more than 15 percentage points in total. The Reads column is that coverage minimum; 23 of the 26 rows are from the one-read search. Percentages are coordinate-level estimates, so Δ can differ from the matrix effect and can fall below the 20-percentage-point selection floor. Δ is paternal minus maternal, in percentage points. *Pabpc1l2b* at chrX:102,056,725, *Pdha1*, and *Erdr1x* lie in the 2 kb upstream of the transcription start; the other sites lie in the gene body.

| Sex | Coordinate | Gene | Pat. % | Mat. % | Δ | Mice | Reads |
|---|---|---|---|---|---|---|---|
| female | chr2:174,140,344 | *Gnas* | 27.0 | 40.1 | −13.1 | 3 vs 4 | 1 |
| female | chr2:174,141,532 | *Gnas* | 27.8 | 52.5 | −24.7 | 3 vs 4 | 1 |
| female | chr2:174,170,967 | *Gnas* | 36.3 | 54.6 | −18.3 | 4 vs 4 | 1 |
| female | chr5:111,567,906 | *Mn1* | 10.4 | 29.2 | −18.8 | 4 vs 4 | 1 |
| female | chr5:143,239,986 | *Zfp316* | 25.4 | 52.6 | −27.2 | 3 vs 4 | 1 |
| female | chr5:143,240,508 | *Zfp316* | 44.3 | 67.1 | −22.8 | 3 vs 4 | 1 |
| female | chr7:125,891,632 | *Sbk1* | 91.7 | 68.3 | +23.4 | 4 vs 3 | 1 |
| female | chrX:102,056,725 | *Pabpc1l2b* | 25.7 | 51.0 | −25.3 | 3 vs 3 | 5 |
| female | chrX:102,057,693 | *Pabpc1l2b* | 31.7 | 58.7 | −27.0 | 4 vs 3 | 5 |
| female | chrX:104,808,131 | *Fgf16* | 45.2 | 24.6 | +20.6 | 4 vs 4 | 1 |
| female | chrX:149,830,719 | *Fgd1* | 16.7 | 41.5 | −24.8 | 4 vs 4 | 1 |
| female | chrX:151,790,673 | *Usp51* | 5.0 | 35.0 | −30.0 | 4 vs 4 | 1 |
| female | chrX:156,826,260 | *Cnksr2* | 43.8 | 20.7 | +23.1 | 4 vs 4 | 1 |
| female | chrX:158,921,595 | *Pdha1* | 11.4 | 28.6 | −17.2 | 3 vs 4 | 1 |
| female | chrX:168,792,821 | *Erdr1x* | 65.4 | 84.4 | −19.0 | 4 vs 4 | 5 |
| female | chrX:169,104,322 | *Akap17a* | 100.0 | 83.3 | +16.7 | 4 vs 3 | 1 |
| female | chrX:169,110,696 | *Asmt* | 66.7 | 100.0 | −33.3 | 3 vs 4 | 1 |
| male | chr2:174,170,876 | *Gnas* | 47.9 | 13.3 | +34.6 | 4 vs 3 | 1 |
| male | chr5:143,239,206 | *Zfp316* | 47.1 | 74.2 | −27.1 | 4 vs 4 | 1 |
| male | chr5:143,240,421 | *Zfp316* | 54.2 | 79.2 | −25.0 | 4 vs 4 | 1 |
| male | chr7:105,411,804 | *Dchs1* | 38.8 | 64.2 | −25.4 | 4 vs 4 | 1 |
| male | chr7:143,013,171 | *Cdkn1c* | 56.7 | 37.8 | +18.9 | 3 vs 4 | 1 |
| male | chr7:143,014,076 | *Cdkn1c* | 52.8 | 29.2 | +23.6 | 3 vs 4 | 1 |
| male | chr10:127,162,287 | *Arhgap9* | 4.2 | 25.0 | −20.8 | 4 vs 4 | 1 |
| male | chr13:119,624,141 | *3110070M22Rik* | 29.2 | 50.0 | −20.8 | 4 vs 4 | 1 |
| male | chr15:100,951,244 | *Fignl2* | 11.7 | 47.5 | −35.8 | 3 vs 4 | 1 |

